# Thyroid and androgen receptor signaling are antagonized by CRYM in prostate cancer

**DOI:** 10.1101/2019.12.19.881151

**Authors:** Osman Aksoy, Jan Pencik, Markus Hartenbach, Ali A. Moazzami, Michaela Schlederer, Theresa Balber, Bismoy Mazumder, Jonathan B Whitchurch, Christopher J. Roberts, Martin Susani, Markus Mitterhauser, Rodrig Marculescu, Gero Kramer, Suzanne D. Turner, Sabrina Hartenbach, Simone Tangermann, Gerda Egger, Heidi A. Neubauer, Richard Moriggl, Zoran Culig, Gregor Hoermann, Marcus Hacker, David M. Heery, Olaf Merkel, Lukas Kenner

## Abstract

Androgen deprivation therapy (ADT) remains a key approach in the treatment of prostate cancer (PCa). However, PCa inevitably relapses and becomes ADT resistant. Besides androgens, there is evidence that thyroid hormone thyroxine (T4) and its active form 3,5,3’-triiodo-L-thyronine (T3) are involved in the progression of PCa. Epidemiologic evidence indicates a higher incidence of PCa in men with elevated thyroid hormone levels. The thyroid hormone binding protein μ-Crystallin (CRYM) mediates intracellular thyroid hormone action by sequestering T3 and blocks its binding to cognate receptors (TRa/TRb) in target tissues. We show in this study that low CRYM expression levels in PCa patient samples are associated with early BCR and poor prognosis. Moreover, we found a disease stage-specific expression of CRYM in PCa. CRYM counteracted thyroid and androgen signaling and blocked intracellular choline uptake. CRYM inversely correlated with [18F]fluoromethylcholine (FMC) levels in PET/MRI imaging of PCa patients. Our data suggest CRYM as a novel antagonist of T3 and androgen-mediated signalling. The role of CRYM could therefore be an essential control mechanism for the prevention of aggressive PCa growth.

**Highlights:**

- Thyroid and androgen hormone driven pathways in prostate cancer (PCa) are antagonized by μ- Crystallin (CRYM).
- [18F]fluoromethylcholine uptake and prognostic values in PCa correlate with CRYM protein levels.
- Reduced CRYM expression predicts early biochemical recurrence (BCR) in PCa patients.

## Introduction

PCa is the most frequently diagnosed cancer in men in the Western world. The course of PCa is largely driven by androgen receptor (AR) and, accordingly, androgen ablation therapy is a cornerstone of current therapies(1, 2). A large proportion of patients with advanced PCa become resistant to ADT resulting in lethal castration resistant prostate cancer (CRPC). CRPC is mainly associated with genetic amplifications, mutations, or other aberration in the AR (3, 4). Besides androgen, the role thyroid hormone thyroxine (T4) and its more active form 3,5,3’-triiodo-L-thyronine (T3) in the progression of PCa has not been comprehensively elucidated.

The thyroid hormones released into circulation are mostly bound to plasma proteins and subsequently transported to the cytosol by thyroid hormone (TH) transporters which have diverse binding affinities such as Transthyretin (TTR) and thyroxine-binding globulin (TBG) (5). The activity of TH within the cell is regulated by (i) cell uptake involving transporters such as MCT8, (ii) metabolization by DIO1 and DIO3, two members of the iodothyronine deiodinase family and (iii) sequestration via binding to other proteins (6) such as the cytoplasmic protein CRYM, which is known to bind to T3 with high-affinity (7). Intracellular thyroid hormone function in the prostate is dependent on CRYM expression levels (8, 9). Previous reports demonstrated that CRYM expression was downregulated in PCa patients who underwent ADT (10) and that expression of CRYM is also downregulated in a PCa xenograft tumor model (11). It was shown that CRYM expression is particularly low in therapy refractory PCa patient biopsies (11). We corroborated this finding and showed that CRYM is responsive to androgens in the MDA PCa 2b cell line (10).

The objective of this study was to define the role of thyroid hormone and its regulation by CRYM in PCa. We hypothesized that TH signaling may play an auxiliary role in PCa progression. Here we show that CRYM expression levels are lower in PCa compared to normal prostate tissue and are reduced further in metastatic disease. Moreover, Kaplan-Meier analysis reveals low CRYM expression as a negative prognostic factor. In PCa cell lines CRYM expression significantly suppresses thyroid and androgen-induced genes. Intracellular [18F]-fluoromethylcholine uptake in PCa was also significantly blocked by CRYM. Importantly, [18F]fluoromethylcholine (FMC) PET/MRI *in vivo* imaging of PCa patients correlates inversely with CRYM levels. Our data propose a novel role for CRYM as a strong antagonist of thyroid and androgen hormone pathways. The important function of CRYM could be a control mechanism for the progression of PCa.

## Results

### Low µ-crystallin (CRYM) expression is a negative prognostic factor and a hallmark of PCa

We assessed CRYM expression levels in malignant and adjacent normal biopsies derived from a large PCa patient cohort by immunohistochemistry (IHC). Decreased protein expression of CRYM was observed in PCa patient samples (n=178) compared to normal prostate glands (n=178), and a further reduction in CRYM expression was observed in metastatic samples (n=17, Figure 1A). Representative IHC images of cytosolic CRYM in normal prostate, PCa and metastases are shown in Figure 1A. These findings were validated through a second independent cohort covering PCa and benign prostate tissue samples (Tuebingen Cohort, Figure 1B). PCa specimens (n=122) had lower CRYM protein levels as compared to benign samples (n=30). Strikingly, low mRNA expression of *CRYM* in metastatic PCa samples compared to primary PCa tissue was observed in multiple larger PCa patient cohorts (Figure 1C, E, F, G, H). Furthermore, *CRYM* expression was significantly higher in non-neoplastic prostate glands compared to primary PCa and to samples from metastasized PCa (Figure 1C) (12). Reduced CRYM expression also correlated with advanced Gleason scores or progressive ‘N’ stages of regional lymph node infiltration (supplementary Figure 1A, B, C). Together, these data indicate that loss of *CRYM* expression in PCa is an indicator for PCa aggressiveness. We analyzed the effect of CRYM expression on biochemical recurrence (BCR)-free survival by Kaplan-Meier analysis in 179 PCa Vienna cohort patients. Low CRYM expression was associated with a reduced time to BCR, indicating that low CRYM expression predicts poor prognosis (Figure 1D). A single patient was also assessed for CRYM and TRβ expression longitudinally within a 4-month follow-up using transurethral resections of PCa. During this period, AR and PSA expression were reduced whereas Ki-67 staining was significantly increased, suggesting progression to a highly aggressive androgen-independent neoplasia. Indeed, we found that CRYM expression was markedly reduced and TRß level increased during progression to androgen-independency in line with our previous findings (supplementary Figure 1D). Taken together, the expression and prognostic value of CRYM reveals its role in regulation of AR and TH signaling in PCa.

**Figure 1.**
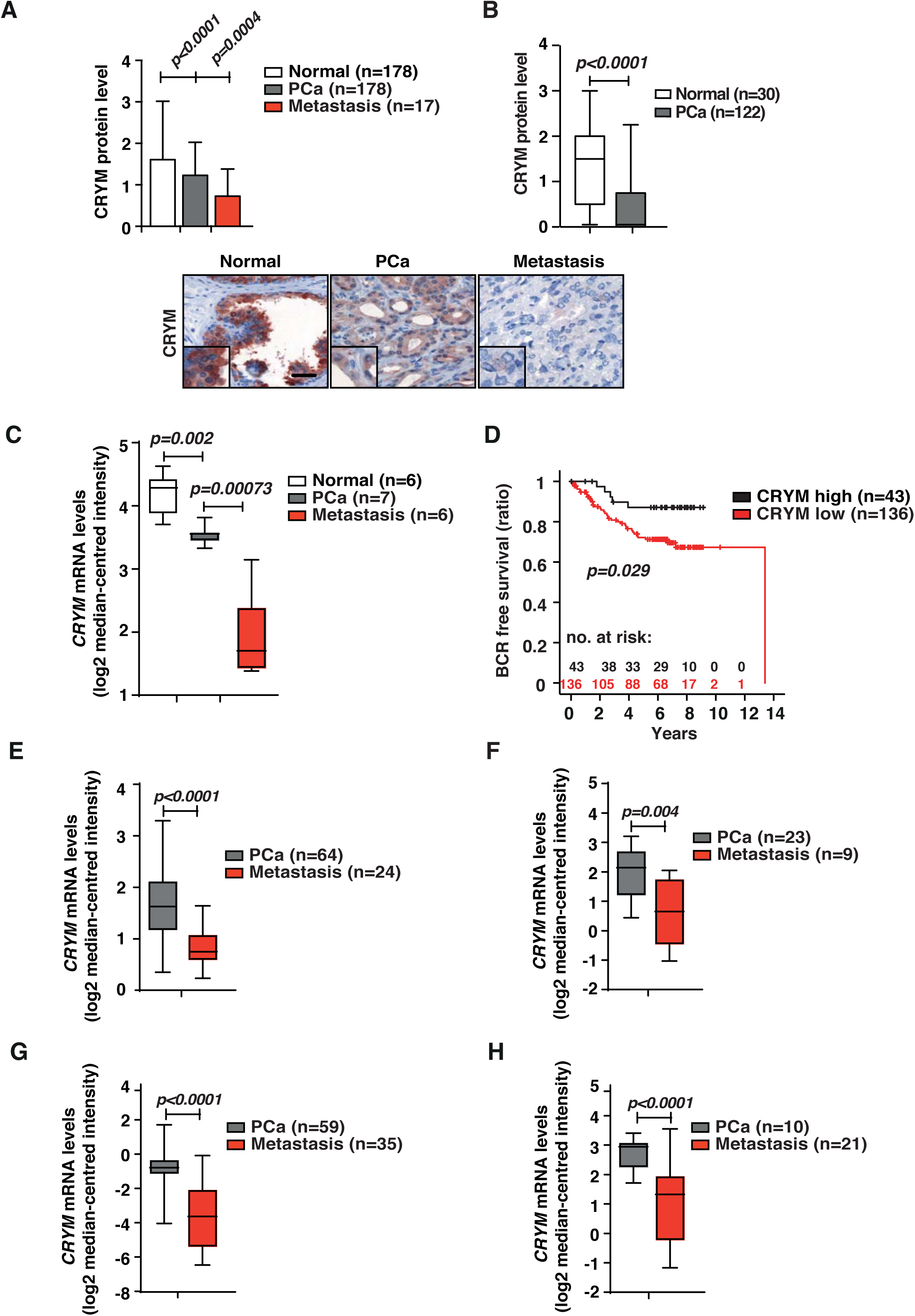
The key binding protein in thyroid hormone metabolism; CRYM is stage- dependently expressed in prostate cancer at RNA and protein levels. **(A)** CRYM levels by IHC of healthy prostate tissue (n=178), PCa (n=178) and tissues derived from PCa metastases (n=17). Staining for CRYM showed significantly reduced levels in healthy tissue versus PCa and further decrease in metastases. Representative staining for CRYM in healthy prostate, PCa and metastatic tissue, scale bar 100µm. **(B)** IHC analysis of CRYM protein levels in non-neoplastic prostate tissue (n=30) and PCa (n=122) in a cohort from Tuebingen. **(C)** The relative expression of CRYM mRNA measured by qRT-PCR in freshly collected tissue samples and paired adjacent non-tumor (Normal n=6, PCa n=7, Metastasis n=6). **(D)** IHC CRYM protein levels in correlation to time for biochemical recurrence (BCR) in a Kaplan-Meier analysis (p=0.029). **(E)** CRYM mRNA expression (Log2 median-centered intensity) is reduced in the patients with metastatic prostate tumors compared to primary tumors (PCa n=64, Metastasis n=24, p<0.001). **(F)** CRYM mRNA expression (Log2 median-centered intensity) is reduced in patients with metastatic prostate tumors compared to that in primary tumors (PCa n=23, Metastasis n=9, p=0.004). **(G)** CRYM mRNA expression (Log2 median-centered intensity) is reduced in patients with metastatic prostate tumors compared to primary tumors (PCa n=59, Metastasis n=35, p<0.001). **(H)** CRYM mRNA expression (Log2 median-centered intensity) is reduced in patients with metastatic prostate tumors compared to that in primary tumors (PCa n=10, Metastasis n=21, p<0.001).

### CRYM enables intracellular accumulation of T3 in PCa cells

To determine the effect of CRYM on intracellular accumulation of T3, we transfected CRYM negative PCa cell lines with empty vector or CRYM expression vector (Figure 2A). Growth medium was supplemented with 10 nM T3 or vehicle, and after 48 hours, T3 levels were measured in the supernatants using a immune-chemiluminescence assay. We observed significantly reduced levels of T3 in the medium of all CRYM re-expressing cells (PC3, 44%; DU-145, 22%; 22RV-1, 20% and LAPC-4, 18% Figure 2A and B) compared to controls. To confirm that this reflected uptake into cells, radioactively labeled thyroid hormone [^125^I] T3 was supplemented to the culture media. More T3 was found in PC3 and LNCaP cell lines with CRYM overexpression compared to empty vector controls (Figure 2C). It can be hypothesized that CRYM binds free T3 in the tumor cells, which results in increased T3 uptake and concomitantly reduced T3 levels in the growth medium.

**Figure 2.**
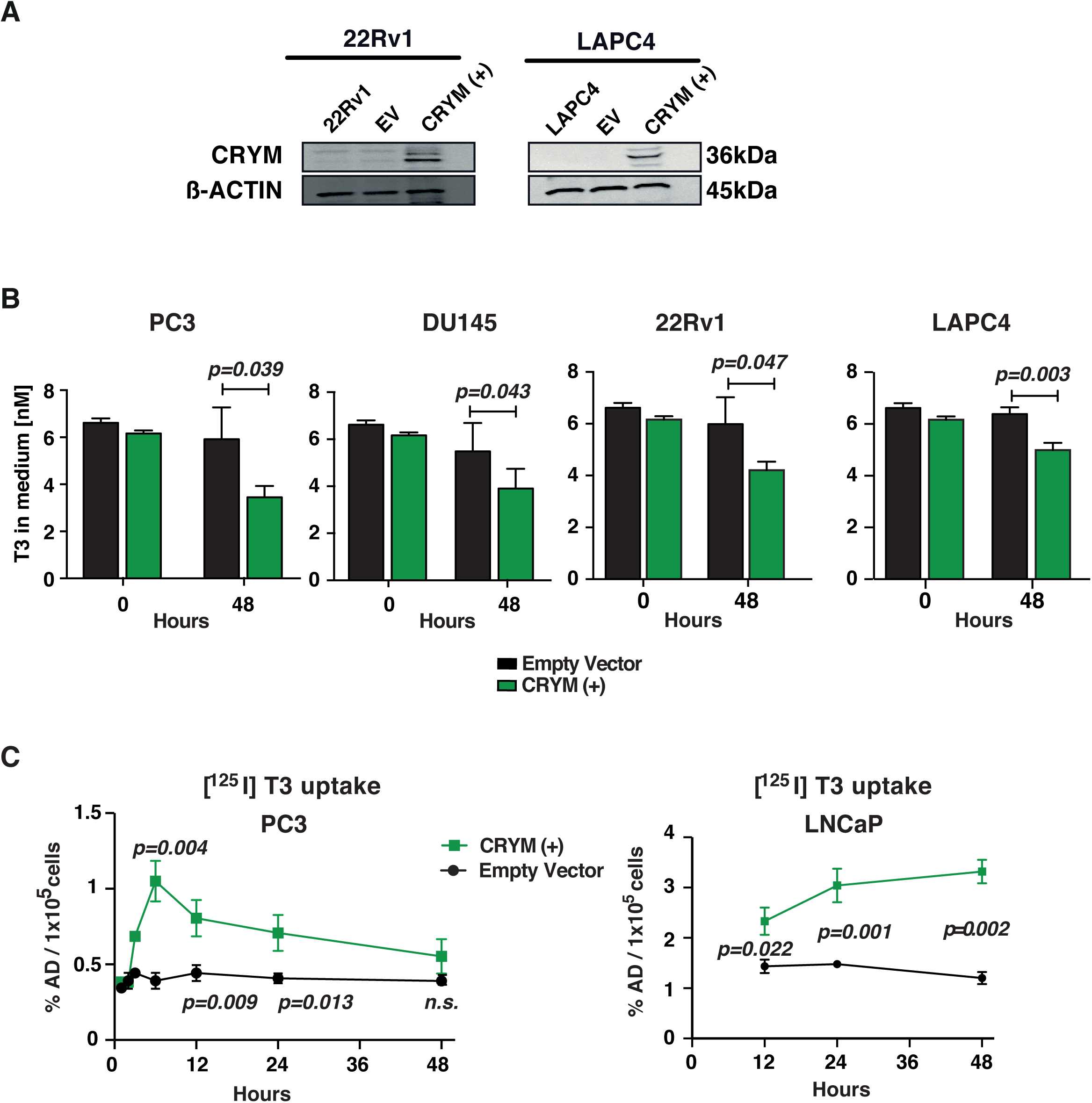
CRYM overexpression leads to a reduction of free T3. **(A)** Cells were transfected with a plasmid bearing the CRYM gene under the CMV promoter that co-expressed GFP. High expression of CRYM was detected in CRYM(+) transfected cell lines 22RV-1 and LAPC-4 by Western blot and was absent in empty vector (EV) and untransfected control. **(B)** T3 concentrations were determined in the growth medium using an electrochemiluminescence immune assay. To test the uptake of T3 in PCa cells we used hormone free medium that was supplemented with a defined amount of T3 (10 nM). At addition of T3 (0) and 48 hours later (48) we measured T3 concentrations in the medium using electrochemiluminescence immune assay. T3 levels were reduced in all cell lines that we tested in medium with CRYM overexpressing cells as compared to control cells bearing an empty vector, suggesting an increased uptake. **(C)** The direct cellular uptake of T3 was quantified. Empty vector and CRYM(+) PC3 and LNCaP cells were incubated for 48 hours in hormone free medium with radioactively labeled T3 ([^125^I] T3). Cells were subsequently washed and intracellular radioactivity was assessed using scintillation counting.

### CRYM reduces the invasive potential of PCa cells by interfering with T3 signaling

WB analysis of CRYM and TRβ expression levels demonstrated that these are mutually exclusive in the androgen-independent PC-3 and DU-145 cell lines. In the androgen-dependent cell lines, both CRYM and TRβ expression could be detected (Figure 3A). Re-introduction of CRYM in PC-3 and DU145 cells reduced TRβ expression (Figure 3B). Next, we monitored the effect of high CRYM expression on invasive potential using matrigel-coated invasion chambers. CRYM overexpression significantly reduced invasion by 50% (p=0.0167) and 67% (p=0.0014) for PC3 and DU-145 cells, respectively (Figure 3C). This is in line with our patient data exhibiting that CRYM protein expression is downregulated in metastasizing tumors whereas TRβ expression is increased. Notably, in two independent PCa patient cohorts CRYM and THRB mRNA levels were inversely correlated in primary PCa and metastasis. Moreover, CRYM expression was significantly downregulated (cohort A: 3,21-fold, *p=0*.*0002;* cohort B: 3,10-fold, *p=0*.*0007)* P and THRB levels concomitantly overexpressed (cohort A: 1,89-fold, *p=0*.*0001;* cohort B: 2,11-fold, *p=0*.*045)* in metastases compared with primary PCa tumor tissue (Figure 3D). Taken together, our data suggest that CRYM binds and sequesters T3 in PCa and correlates with reduced invasive capacity. Since free T3 is known to influence TRβ expression(13), we hypothesize an autoregulatory loop where CRYM masks free T3 and therefore leads to reduced TRβ expression.

**Figure 3.**
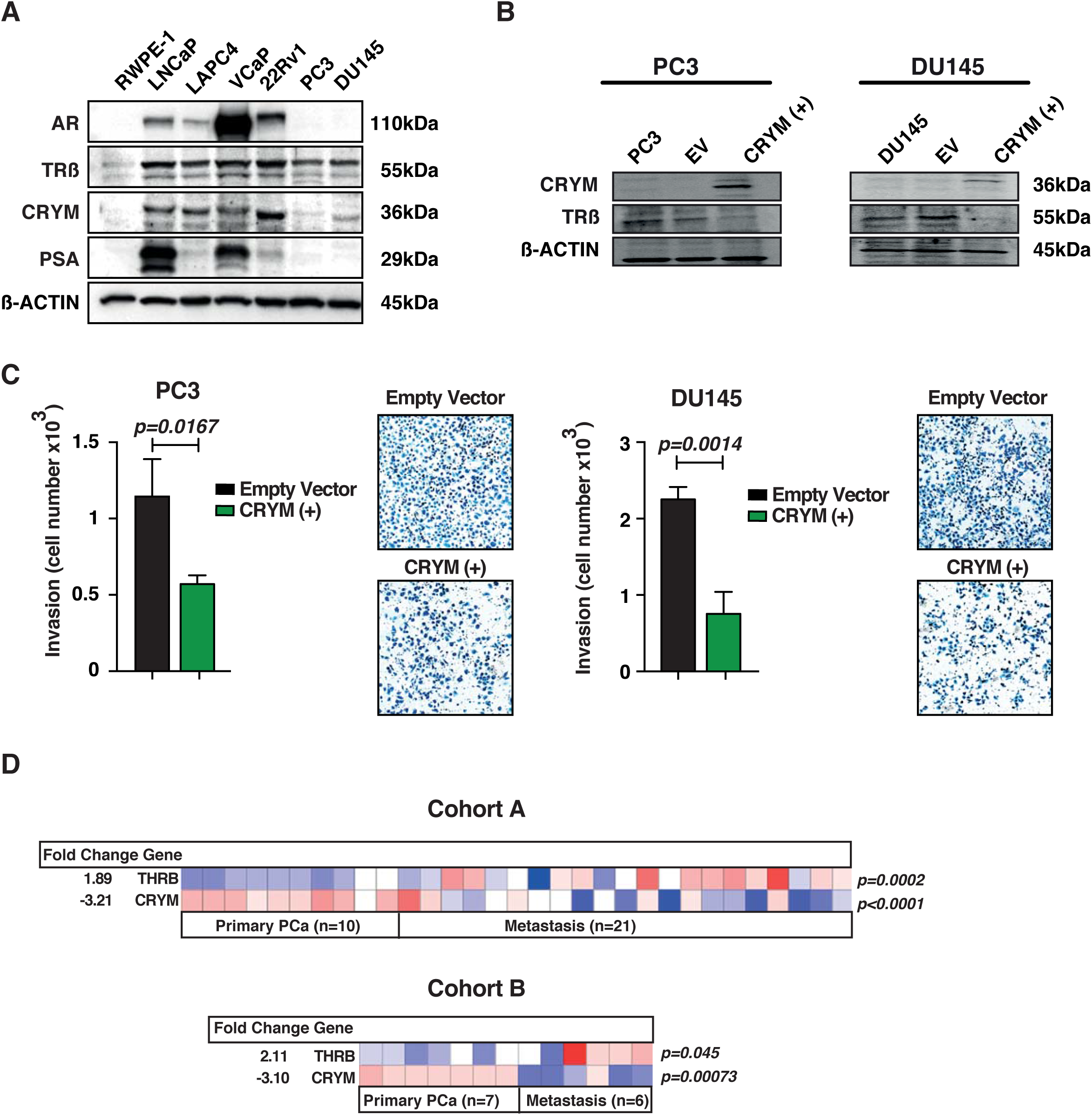
CRYM induced the invasion through repression of T3 signaling. **(A)** Western blot of CRYM and TRβ in PCa cell lines LNCaP, PC3, DU-145, LAPC4 and 22Rv1. Androgen independent, metastasis derived cell lines PC3 and DU-145 did not express CRYM, but were positive for TRβ. The RPWE-1 cell line is derived from normal prostate tissue was negative for CRYM and showed very low TRß levels. The androgen responsive cell lines LNCaP, LAPC4, VCaP and 22Rv1 showed CRYM and TRß expression, β-actin was used as loading control. **(B)** High CRYM expression in the most aggressive PC3 and DU145 led to a reduction of TRβ protein levels. **(C)** Matrigel-coated invasion chambers were used to test invasive capacity of PC cell lines PC3 and DU145 with and without CRYM overexpression against a gradient from 10% to 15% FCS in 1680 RPMI medium for 24 hours. Lower invasive capacity was observed in cells that overexpressed CRYM. **(D)** Diagram of CRYM and TRβ expression fold change in metastatic and primary PCa in two different datasets. CRYM and TRβ expression levels are inversely correlated in primary tumors and metastases. **Cohort A**: CRYM expression is high in primary prostate cancer (n=10) and low in metastases (n=21); TRβ is low in primary prostate cancer (n=10) and high in metastasis (n=21). p<0.0001). **Cohort B**: CRYM expression is high in primary prostate cancer (n=7) and low in metastases (n=6); TRβ is low in primary prostate cancer (n=7) and high in metastasis (n=6). p<0.0001).

### CRYM re-expression changes the transcriptome revealing thyroid and androgen receptor antagonism

Given the proposed antagonistic role of CRYM on thyroid hormone signaling and its negative association with advanced PCa (9, 11), we assessed the impact of CRYM overexpression on gene expression in androgen sensitive and AR expressing LNCaP cells using polyA enriched RNA-sequencing (RNA-Seq). Cells were transfected with a plasmid conferring neomycin resistance containing the CRYM ORF in addition to a GFP under the CMV promoter or empty vector control and were selected in G-418 containing medium. Exogenous CRYM resulted in differential expression of 9.25% of genes (n=3226; 2.72% up, 6.53% down) (Figure 4A). Ingenuity pathway analysis (IPA) was performed on significantly deregulated genes (>2-fold; q<0.05; n=642) to identify over-represented pathways. Selected genes known to be activated by the TR/RXR (n=33) axis were suppressed upon CRYM overexpression including B-cell leukemia 3 (*BCL3*) and fatty acid synthase (*FASN*). High *FASN* expression is a known feature of aggressive PCa and it has been proposed as an metabolic oncogene (14) (15, 16).

**Figure 4.**
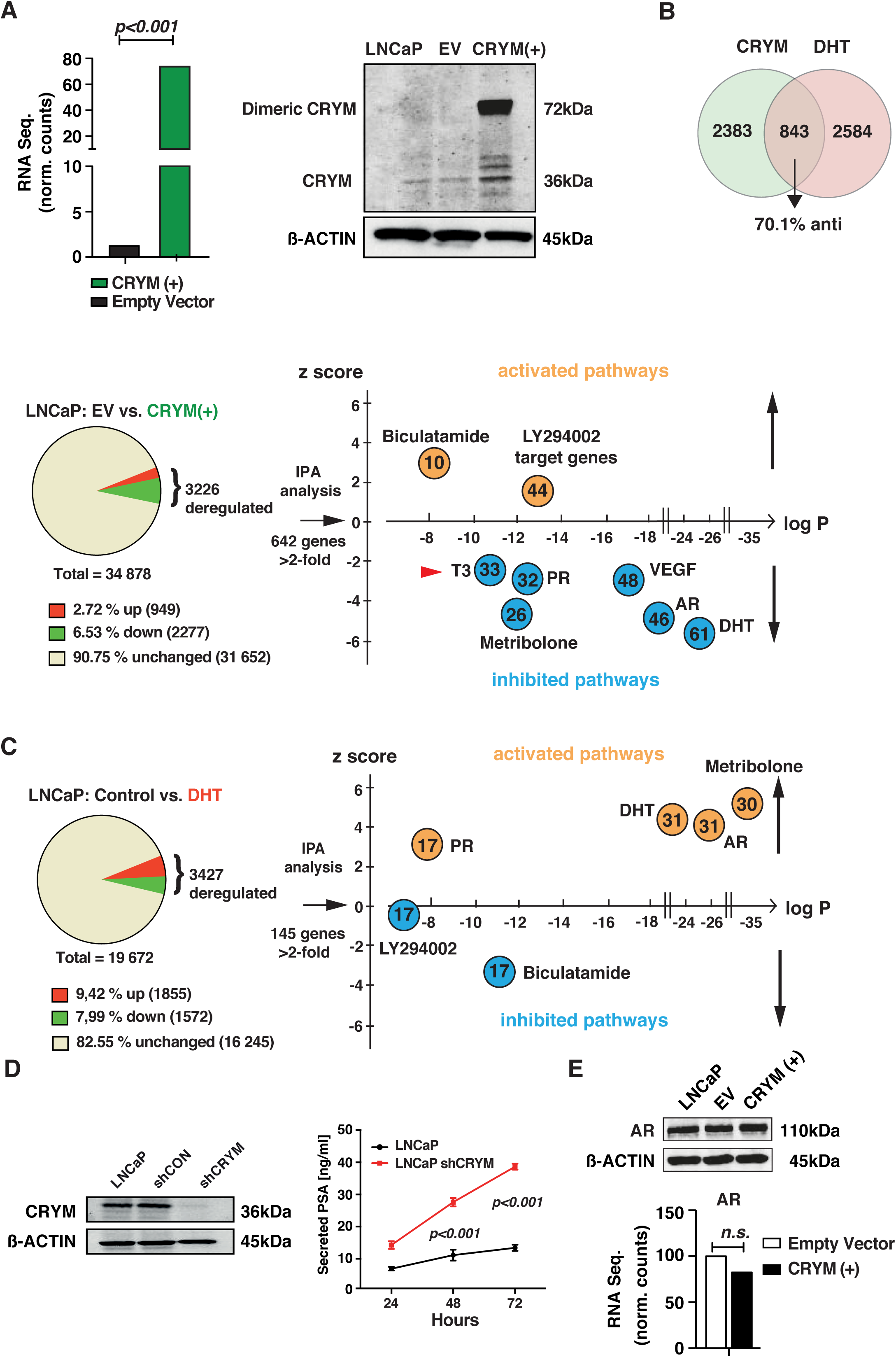
CRYM-induced suppression of T3 and AR signaling. **(A)** Overexpression of CRYM in LnCAP is confirmed by RNA-seq and western blot analysis. We performed single end 50 bp RNA-seq with an Illumina Hi-Seq2000 platform and compared LNCaP CRYM(+) to empty vector cells using two biological replicates for each group. Strong overexpression of CRYM was validated by RNA-seq (73.7 versus vs. 2.2 normalized counts, p*<0*.*001*). **(B)** Of 58,540 assessed RNA types 34,878 (59.6%) were expressed in LNCaP cells. CRYM overexpression led to significant deregulation in 9.25% of these (2.72% up, 6.53% down; >2-fold). Significantly deregulated genes (*p<0*.*05*, >2-fold, n=642) were analyzed using the IPA analysis to identify upstream regulators of deregulated genes (enrichment of target genes as reflected in *p*-value) and activation or inhibition of the respective pathway, reflected in a positive or negative z score, respectively. We found highly significant enrichment and inhibition of pathways involved in androgen signaling, indicating a tight interaction of thyroid and androgen signaling (circles represent target genes of the respective pathway: Progesterone receptor (PR), Vascular endothelial growth factor (VEGF), androgen receptor (AR), dihydrotestosterone (DHT). **(B and C)** Analysis of a published data set of genes deregulated by androgen DHT in the same cell line and compared it to our data (17). We found that 843 genes overlapped amongst CRYM and DHT regulated genes. 70.1% of these were counter-regulated (anti), which suggests partly antagonistic functions of CRYM and DHT. **(D)** CRYM knockdown in the LNCaP cell line was generated using lentiviral-transfected shRNAs (shCRYM) and a non-targeting control shRNA (shControl). CRYM reduction was confirmed by Western blot. PSA release levels were measured by a chemiluminescence immune assay. Cells with CRYM knock-down had consistently higher PSAs in the growth medium. **(E)** AR expression was tested in association with CRYM overexpression at protein (Western blot) as well as mRNA (RNA-seq) levels resulting in no changes, thus excluding a direct effect of CRYM on AR expression.

Interestingly, genes involved in AR signaling including dihydrotestosterone (DHT; n=61) and AR regulated (AR; n=46) genes were significantly downregulated (Figure 4A). We then compared our RNA-Seq dataset of CRYM overexpressing cells to a published RNA-Seq. dataset(17) wherein the same cell line had been treated with DHT (Figure 4B and C). Interestingly, among the overlapping, deregulated genes, 70.1% were counter-regulated, suggesting antagonistic functions for CRYM and AR signaling. In this dataset, 843 out of 3427 significantly deregulated genes overlapped with the CRYM deregulated genes (Figure 4C). We performed IPA analysis on the DHT deregulated genes (>2-fold, q<0.05; n=145). As expected, progesterone, DHT, AR and metribolone regulated genes were strongly enriched and the respective pathways activated, whereas Bicalutamide and LY294002 associated genes were downregulated (Figure 4C). Complementary to this approach, shRNA-mediated knock-down of CRYM in LNCaP cells resulted in a significant increase in PSA levels (38±2ng/ml as compared to 10±2 ng/ml of control, Figure 4D). Since AR protein or mRNA levels were not altered, the strong effect of CRYM overexpression on target gene expression, suggests a crosstalk between androgen and thyroid hormone signaling in PCa (Figure 4E).

### CRYM alters the choline metabolism and metabolic shift caused by T3

Recent studies have revealed that tumor suppressor pathways such as PTEN, p53 and RB are involved in regulation of glucose and glycine metabolism resulting in metabolic features that are unique for proliferative cancer cells. We took advantage of the 1H-NMR-based metabolomics to identify the role of CRYM in thyroid hormone signaling and metabolism during PCa progression. PC3 harboring empty vector or CRYM overexpression were stimulated with T3 for 48 hours. We performed intracellular and extracellular metabolite concentrations analysis followed with principal component analysis and found that metabolomic profile of T3 or CRYM overexpression has significantly affected PCa cells (Figure 5A), but did not lead to profound change in the metabolomic profile of non-transformed cells (data not shown). CRYM overexpression had a strong overall effect on the metabolome (represented by a shift along the t(1) axis in partial least squares-discriminant analysis (PLS-DA) regardless of T3 stimulation. Our NMR metabolomic analysis showed that intracellular choline was significantly reduced upon CRYM overexpression in PC3 cells, whereas T3 led to a slight but non-significant choline increase (Figure 5B). Choline is an important component for phospholipid metabolism (17) and elevated choline is a metabolic hallmark of tumor progression and a valuable diagnostic biomarker in PCa (18) (19). Next we tested the effect of T3 on choline kinase α (CHKA) expression. Phoshorylation of choline by this enzyme is the first step of phosphadidylcholine synthesis. We found that T3 supplementation increased CHKA expression. *Vice versa* we described above that CRYM suppresses fatty acid synthase (FASN) which is another essential component for membrane phospholipid synthesis. Of note, a recent report showed that CRYM knockout mice have increased body weight and PPARγ expression under high fat diet adding a new aspect on the connection of CRYM and lipid metabolism (20)

**Figure 5.**
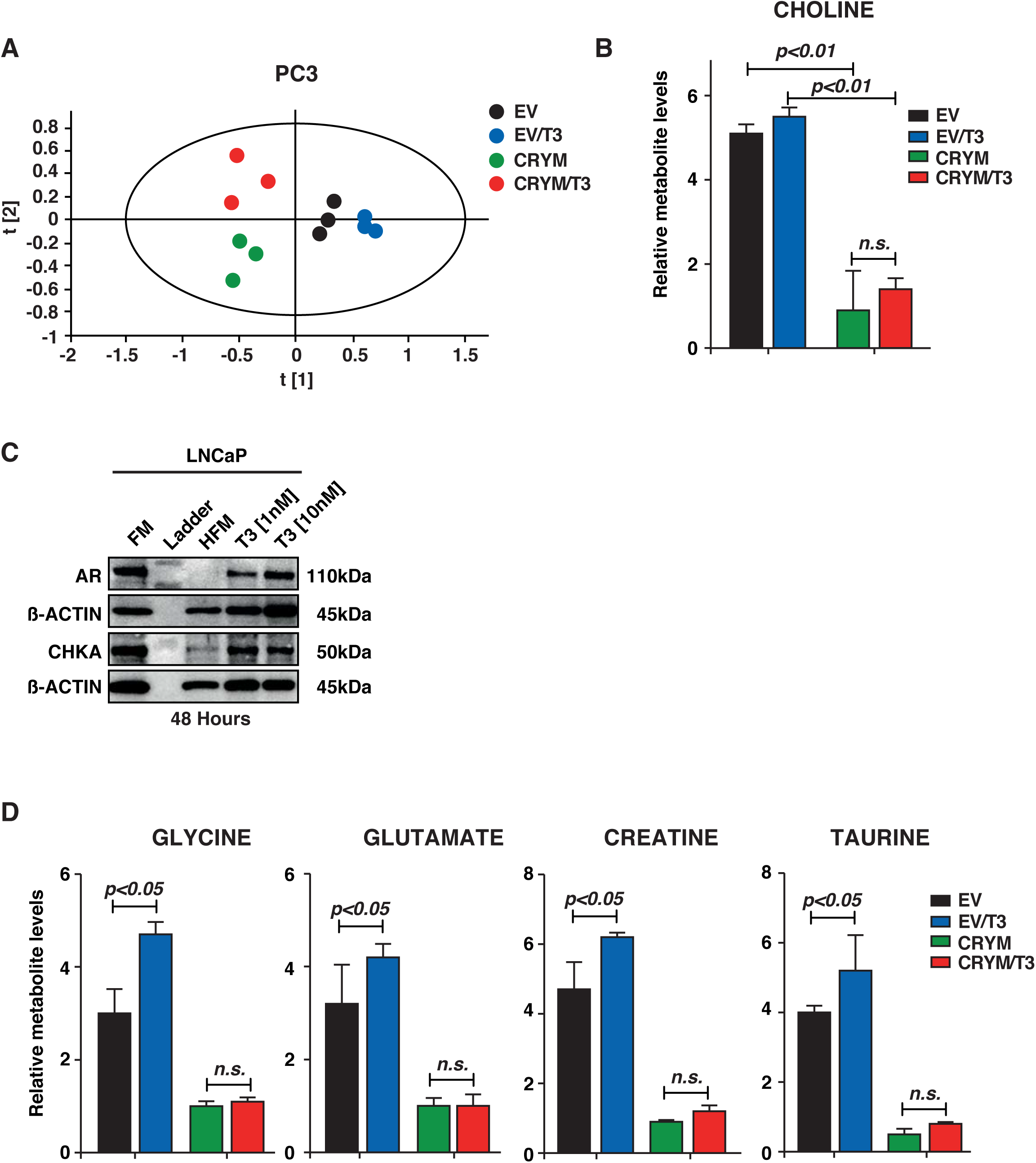
CRYM overexpression caused to the metabolic alterations in PCa. **(A)** Score plot of partial least squares-discriminant analysis (PLS-DA) model of PC3 cell pellet extract fitted using 1H-NMR spectral data. PC3 cells without CRYM overexpressing vector (right) were separated from PC3 cells with CRYM overexpressing vector (left) along the first component. **(B)** Free choline was drastically reduced by CRYM overexpression (p<0.01) irrespective of the presence of T3. **(C)** Choline kinase α (CHKA) as an androgen receptor chaperone in RPMI 1640 medium with 10% FCS (FM), HFM and HFM with T3 supplementation (1 nM and 10 nM). T3 leads to stabilization of the AR receptor by inducing AR chaperone choline kinase *α* (CHKA). CHKA was expressed very low under HFM conditions and could be rescued by T3 supplementation. **(D)** Relative metabolite levels of glycine, glutamate, creatinine and taurine were significantly induced by addition of T3 and reduced by overexpression of CRYM regardless of T3 as measured by ^1^H-NMR.

Glycine, glutamate, creatine and taurine, which are known to be rapidly taken up by growing cancer cells, were increased by T3 but reduced with CRYM overexpression, reflecting the inhibition of cancer associated metabolism under CRYM overexpression and suggesting that supporting CRYM expression can abolish the stimulating metabolic influence of T3 on PCa cells (Figure 5D).

### FMC-PET [18F]fluoromethylcholine imaging of PCa patients is a surrogate marker for thyroid hormone action in PCa

Radiolabeled choline such as [18F]fluoromethylcholine (FMC) is widely used as PET/MRI tracer for the evaluation of PCa progression. This is potentially due to the increased rate of lipid metabolism in neoplastic prostate tissues where choline is needed to generate phosphatidylcholine, an important component of the cell membrane (21). Having in mind the effect of CRYM on intracellular choline metabolism in PC3 PCa cells, CRYM and TRβ protein expression by IHC were analyzed in the same patient prostatectomy specimens. Figure 6A shows whole mount sections of two different PCa specimens along with the respective FMC-PET results. A representative tumor with a Gleason pattern 3 (GL3) lesion displayed a lower FMC signal along with high CRYM-low TRβ expression (left panel) compared to a Gleason pattern 4 (GL4) PCa lesion with higher FMC signal and low CRYM-high TRβ expression (right panel). Next we measured FMC levels in a cohort of 42 PCa patients before they underwent radical prostatectomy. Statistical evaluation of 42 patients confirmed a direct correlation for PET-FMC uptake signal to IHC TRβ levels and an inverse correlation to IHC CRYM levels in the respective tumor tissue (Figure 6B). A follow up analysis was performed in 87 patients that correlated with FMC uptake to biochemical recurrence and/or synchronous metastatic disease. Mean follow up time in this cohort was 508 days. Patients with biochemical recurrence or already initial metastases had significantly higher FMC uptake (Figure 6C). ROC analysis shows the confidence interval curve of 0.77 with p<0.0001 (Figure 6D). These data indicate that FMC signal in prostate is indicative of BCR, and that this correlates with TRβ and CRYM expression. Our results suggest that choline is closely associated with intracellular thyroid hormone levels and that the ^18^F-FMC PET/MRI tracer for the activity of thyroid hormone could be used to predict high and low risks in PCa patients. Whether FMC-PET imaging can be used as a marker for the activity of thyroid hormone metabolism in PCa needs to be tested in further studies. In summary we conclude that absence of CRYM results in an increased choline metabolism in PCa, which is a poor prognostic indicator that can non-invasively be measured in vivo by ^18^F-FMC PET/MRI.

**Figure 6.**
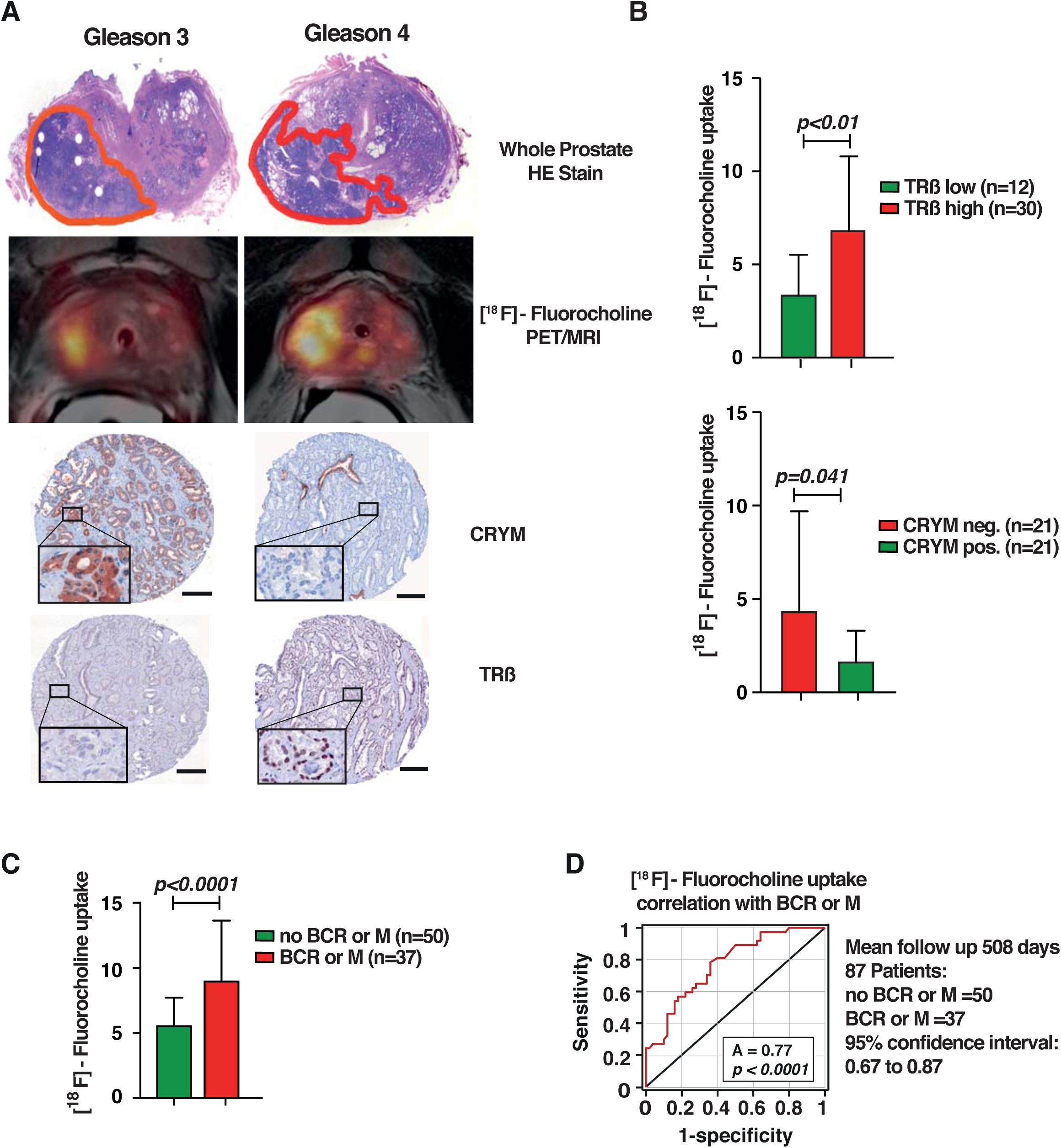
Non-invasive imaging using [^18^F]fluoromethylcholine (FMC) and positron emission tomography (PET) is a surrogate marker for the activity of thyroid hormone metabolism in PCa. **(A)** H&E whole mount sections of two different prostate cancer specimens on the left with mainly Gleason 3 lesion and on the right with larger Gleason 4 lesion. Corresponding FMC PET/MRI of the same patient shows low FMC uptake on left (Gleason 3) and high FMC uptake on the right (Gleason 4). IHC shows intensive CRYM staining and almost no TRβ levels on the less aggressive left side but high TRß on the more aggressive right side, on which CRYM is low. **(B)** Statistical evaluation of 42 PCa patients that had FMC PET/MRI before radical prostatectomy. CRYM and TRβ protein levels in tumors were analyzed using IHC. Choline uptake was significantly increased in tumor specimens with high TRβ (TRβ 2-6 score; n=30, p<0.01) or no CRYM expression (n=21, p=0.041). **(C)** Choline uptake correlated to BCR or metastases in receiver-operating-characteristics-curve analysis (AUC=0.77, p<0.0001). **(D)** 87 Patients with FMC PET/MRI and a mean follow-up time of 508 days were divided into a group that developed BCR and/or metastasis (n=37) and a group that did not (n=50). The first group had significantly higher choline levels in FMC PET/MRI (p<0.0001).

## Discussion

We investigated the crosstalk of CRYM with T3 and AR signalling in PCa. We demonstrated that the growth-promoting effect of T3 could be attenuated by the intracellular thyroid hormone binding protein CRYM. Increased CRYM expression resulted in decreased availability of T3, significantly prolonged time to BCR and suppressed important targets of the androgen-regulated gene network in PCa. Loss of CRYM in aggressive PCa cells led to a massive increase in PSA. We show here for the first time a tumor suppressive effect of CRYM by reducing choline metabolism in PCa. In FMC-PET/MRT studies of PCa patients we found a significant correlation of high CRYM expression with low choline uptake. Expression of CRYM may be a novel biomarker for the diagnosis and prognosis of PCa. We propose that targeting the thyroid hormone pathway is a rational strategy to treat aggressive PCa.

Besides androgens, some other factors such as thyroid or estrogen hormones have been proposed to take part in the progression of PCa (22, 23). Recent epidemiological studies have linked high thyroid hormone levels to higher incidence of PCa suggesting a tumor promoting role of T3 in PCa (24). However, underlying molecular mechanisms in which thyroid hormone signaling contributes to tumor growth are still not well understood.

The growth-promoting effect of T3 can be attenuated by the intracellular thyroid hormone binding protein CRYM. Increased levels of CRYM lead to suppression of important targets of the androgen-regulated gene network including PSA. We found high CRYM levels in normal human prostate samples, decreased CRYM in PCa samples and CRYM expression couldn’t be detected in metastatic PCa. Patients with high CRYM levels in their PCa cells had a significantly prolonged time to BCR, suggesting CRYM as a positive predictive marker in PCa. Since CRYM is involved in sequestration of T3, the loss of thyroid hormone binding protein CRYM in PCa could result in increased thyroid hormone activity. Our data on CRYM/T3 regulation further support the idea that CRYM acts not only as a binding protein but also as a T3 sink in the cytoplasm. CRYM overexpression downregulated thyroid and androgen regulated genes which indicates a possible interaction of thyroid and androgen signaling pathways in PCa. High CRYM levels could possibly antagonize the impact of thyroid hormone on androgen signaling.

PET imaging using metabolic tracers offers the opportunity not only for non-invasive diagnostics but also long-term surveillance. Interestingly, we found that overexpression of T3-buffering CRYM leads to a reduction in intracellular choline levels in PC3 cells. CHKA has a crucial role in phospholipid metabolism through catalyzing the choline phosphorylation to form phosphocholine, a phospholipid component of bilayers in cell membranes (25). Total choline expression has been shown to be elevated in PCa tissue as compared to healthy, age matched controls (26-28). This feature has been successfully used in combination with PET/MRI to enhance the accuracy of non-invasive diagnostics (29). The clinical outcome of PCa is highly variable reflecting the genetic and pathophysiologic heterogeneity of the disease. Therefore, biomarkers for detection of aggressive tumors are highly warranted. The results of our study contribute to define aggressive PCa cases and suggest a novel targeting approach for CRPC. Of note, expression of CRYM could be applied not only as prognostic but also as biomarker to guide patient care and management. Our FMC-PET/MRT cohort encompassed 42 patients displaying a significant correlation with CRYM and also TR*β* status. It is clear that larger, independent patient series are needed before clinical applicability of our findings in the context of FMC-PET/MRT. In addition, PCa screening based on serum PSA measurements shows limited sensitivity and specificity. Thus, it remains to be determined if FMC-PET/MRT could be used to non-invasively discriminate early stage PCa from non-malignant conditions associated with serum PSA increase. In addition, our data provide now a mechanistic rationale for this diagnostic technique, suggesting that choline metabolism is tightly linked to intracellular thyroid hormone status. For treatment of malignancies Propylthiouracil (PTU), a thyroid hormone synthesis blocker that generates a hypothyroid state was shown to reduce tumor growth of engrafted lung and PCa cells in NCR-Mice (7). Hypothyroidism was shown to slow down growth of aggressive breast cancer (30). Earlier reports indicated also that hypothyroidism induced by PTU in rats inhibited metastatic growth, formed smaller tumors and prolonged survival in liver cancer(31). The question of how hypothyroidism diminishes growth of PCa could be possibly addressed in the future with more specific drugs with less side effects blocking the thyroid hormone pathway that could be used in the combination with AR blockers (32, 33).

In conclusion, we provide strong evidence for the protective effect of T3-buffering by CRYM in PCa. Therefore, CRYM might represent a novel biomarker for detection of good prognostic PCa. In addition, we provide a rational for how thyroid hormone metabolism could influence choline levels in PCa, which is important not only for diagnostic imaging techniques. Based on the fact that thyroid hormone signaling might act as a new oncogenic factor, our study also contributes to the understanding of aggressive PCa. Importantly new treatment avenues using novel and specific anti-thyroid drugs may open up and inclusion of such drugs in currently used regimens for metastatic PCa will be of interest.

## Materials and methods

### Cell lines

PCa cell lines LNCaP, PC3, DU145, 22Rv1 were obtained from ATCC (Manassas, VA, USA) and cultured in RPMI 1640 medium (Gibco® Life Technologies, Carlsbad, CA) supplemented with 10% FCS (Gibco®) and 1% penicillin/streptomycin (Gibco®) at 37°C in 5% CO_2_ in a humidified chamber. For hormone sensitive experiments, charcoal stripped FCS was used (CS-FCS, Gibco® Life Technologies). The human PCa cell line LAPC4 was kindly provided by Zoran Culig (Innsbruck Medical University) and cultured in Iscove’s Modified Dulbecco’s Medium (Sigma-Aldrich, St. Louis, MO) supplemented with 10% FCS, 1% penicillin/streptomycin. RWPE-1 (ATCC) was grown in Keratinocyte serum-free media with L-glutamine (Invitrogen) supplemented with 2.5 mg EGF (Invitrogen) and 25 mg Bovine Pituitary Extract (Invitrogen). All cell lines were assessed to be free of mycoplasma infection and were passaged using trypsin/EDTA (Gibco®).

### Prostate cancer immunohistochemistry and human tissue microarray

FFPE samples were obtained from patients that underwent radical prostatectomy (Department of Pathology, Medical University Vienna, Austria, and Institute of Pathology, Tübingen, Germany). Immunohistochemistry (IHC) was carried out according to standard protocols and protein expression quantified as described (Schlederer et al., 2014). The following antibodies were used in this study: CRYM (1:100, Abnova), TRβ (1:100, Rockland), AR (1:250, DAKO), Ki-67 (Leica Microsystems), PSA (1:20 Dako), Cleaved Caspase 3 (CC3) (1:200 dilution; Cell Signaling). TMAs were evaluated by four board certified pathologists (MS, PM, AH, LK) and statistical analysis was performed as described(34).All human samples for TMA and their use in this study were approved by the Research Ethics Committee of the Medical University Vienna, Austria (1248/2015) and by the Research Ethics Committee for Germany (395/2008BO1) (Bonn, Germany). Biochemical Recurrence (BCR) was defined as PSA progression with ≥ 25% increase in PSA and PSA levels above 0.2 ng/ml. For BCR-free survival analysis, Kaplan Meier survival curves and Log-Rank test were conducted. Following R/Biocondor packages were used: survival, survminer. TMAs were performed on the respective tumor areas as well as BPH areas. After IHC staining for CRYM and TRβ as described above, TMAs were assessed by 3 pathologists (S.H., S.R., L.K.) according to 4 points scale for staining intensity (0 to 3) and a 4-point scale for the number of stained cells (0 to 3). A sum score (0 to 6) served as the reference for correlation with semiquantitative FMC-PET data. Patients received a routine clinical follow-up monitoring using PSA serum blood sampling. Biochemical relapse was defined as rising PSA levels above 0.2 ng/ml. For initial staging, synchronous metastatic disease was defined by positive lymph node findings after surgery or biopsy of imaging positive distant lesions.

### Analysis of CRYM and THRB mRNA expression in human tumor samples

For the PCa gene expression cohort, 248 localized/locally advanced PCA patients commencing radical radiotherapy (with/without ADT) at the Northern Ireland Cancer Centre, Belfast Health and Social Care Trust (BHSCT), between 1 January 2005 and 31 December 2009 were included (Jain et al., 2018). Patients were treated with 70– 74 Gy external beam radiation therapy (EBRT) in 2 Gy fractions with 3D conformal or intensity modulated techniques over 7–7.5 weeks. Node negative patients received elective pelvic nodal irradiation at the physician’s discretion; node-positive patients had radiotherapy to pelvic nodal regions. Short (6months) or long (>6–36months) course ADT commenced at least 3months before radiation with LHRH agonists or antiandrogens. Ethical approval for this study was obtained from the Northern Ireland Biobank (NIB Reference 15-0169) under the remit of the NIB’s ethical approval from the Office of Research Ethics Committees Northern Ireland for the collection, storage and release of tissue (ORECNI Reference 16/NI/0030).

### Lentiviral Transduction

To establish CRYM knock down LNCaP cell lines, cells were transduced with the Lentiviral particles targeting the human CRYM coding sequence (Mission shRNAlentiviral particles, Sigma) and a non-targeting shRNA construct as a control using a spinoculation procedure. Briefly, 1.5 × 10^4^ of LNCaP cells were seeded in a 96-well plate and incubated in a humidified chamber at 37°C and 5% CO_2_ to accommodate a confluency of 70-80% on the day of transduction. The medium was removed and replaced with 110 μl fresh medium containing 8 μg/ml Polybrene (Sigma) added to each well to increase the transduction efficiency. A quantity of 10 μl of lentiviral particles was added to the appropriate wells and the cells and lentiviral particles containing plates were immediately centrifuged at 32°C for 90 minutes at 2500 rpm speed in a centrifuge (Beckman Allegra X-12R). After spinoculation, plates were incubated for 24-48 hours at 37°C in a humidified chamber and cells were washed with PBS to remove residual lentiviral particles. Stable clones of LNCaP cells were selected by adding 1 ug/ml of puromycin antibiotic (Millipore).

### Colony formation

To examine the effect of T3 on colony formation, a bottom layer containing RPMI 1640 medium (10% CS-FCS) and 0.75% low melting agar (Bio-Rad) was overlain with half the volume of RPMI 1640 medium (10% FCS) and 0.36% agar containing 3×10^4^ cells. Finally, the two layers were covered with liquid RPMI 1640 medium containing 10% CS-FCS (Gibco® Life Technologies). The liquid top layer with or without 100 nM T3 was changed every 3 days. Cells were incubated at 37°C for 14 days. The top layer was discarded and the 2 remaining layers were flushed with phosphate buffered saline (PBS). Colonies were fixed and stained with crystal violet. The colonies were counted and photographed (Nikon D90).

### Quantification of PSA and T3

Lentivirally transduced LNCaP cells were seeded at a density of 10^5^ cells/ml in 6-well plates and were cultured in RPMI 1640 with 10% FCS for 48 hours. The culture medium was removed and replaced with RPMI 1640 medium with 10% CS-FCS (Gibco® Life Technologies) and cells were incubated with or without 50 nM T3 for another 3 days. The growth medium was collected at 24, 48 and 72 hours. For T3 analysis, PCa cells were transfected by lipofection with empty vector or CRYM expressing vector. After 48 hours, medium was replaced with CS-FCS medium with or without T3. The culture medium was collected at 24 and 48 hours. The level of secreted prostate specific antigen (PSA) and T3 in the growth medium was determined using a validated Elecsys electrochemiluminescence immune assay (Roche, Rotkreuz, Switzerland) on a Cobas e601system (Roche) that is routinely used for clinical applications.

### Western blotting

Cells were lysed using radioimmunoprecipitation assay (RIPA) buffer supplemented with a cocktail of protease inhibitors (Roche). Western blot analysis was performed using standard procedures. Protein lysates were separated by SDS-polyacrylamide gel electrophoresis and then transferred to polyvinylidene difluoride (PVDF) membrane. The membranes were blocked with 5% w/v non-fat dry milk and incubated with CRYM (M03 clone 6B3, Abnova) and TRβ (clone 2386 IgG1, Rockland), AR (Santa Cruz Biotechnology), PSA (Cell Signaling Technology), Choline Kinase α (Chkα, Cell Signaling) antibodies at a dilution of 1:1000. β-Actin and β-tubulin (1:10000 dilution, Cell Signaling Technology). Secondary mouse (315-0035-008) and rabbit (111-036-047) antibodies were obtained from Jackson Immune Research.

### RNA and qPCR

Total RNA was isolated from cultured cells homogenised in QIAzol Lysis Reagent (Qiagen, Germany) using a single-step guanidine isothiocyanate/phenol/chloroform-based extraction technique. RNA concentration and quality were measured using a NanoDrop® 2000 UV-Vis Spectrometer (Thermo Fisher Scientific, Wilmington, USA). Using RevertAid First Strand cDNA Synthesis Kit (ThermoFisher Scientific) 1,000 ng of RNA were reverse-transcribed to cDNA. For qPCR analysis CFX96 Real-Time PCR Detection System (BioRad, Hercules, CA, USA) was employed using Kapa Sybr Fast qPCR Master Mix (2x) Kit (Kapa Biosystems, Wilmington, MA, USA). RNA expression of target genes relative to GAPDH was quantified by 2ΔΔCT-method. Statistical analysis was performed in GraphPad Prism 6.

### Wound healing and invasion assay

Cells were grown in 24-well plates with cell culture inserts for wound healing assays. After 48 hours, medium and inserts were removed to create a wound/gap. New medium with 10% CS-FCS (12676-011, Gibco) with or without 100 nM T3 was added and cells were incubated for a further 2 days. Migration into the empty gap was monitored using a microscope equipped with a camera. For invasion assays, matrigel-coated invasion chambers (BD Biosciences, NJ, USA) were rehydrated for 2 hours in RPMI 1640 medium (Gibco® Life Technologies) supplemented with 10% FCS (Gibco®). Cells (2.5×10^4^/ml) were seeded in the upper chamber in RPMI 1640/10% FCS, whereas the lower chamber contained RPMI 1680/15% FCS. After 24 hours at 37°C, medium was discarded and non-invaded cells were removed with cotton swabs from the top of the membrane, cells that migrated to the bottom of the membrane were fixed with methanol and stained with Toluidine blue. Invading cells were counted using a microscope equipped with a camera.

### Gene expression profiling

Total RNA was extracted using TRIzol® Reagent (Ambion, Thermo Scientific, CA, USA). RNA-Seq libraries were prepared with a TruSeq Stranded mRNA LT sample preparation kit (Illumina) using Sciclone and Zephyr liquid handling robotics (PerkinElmer). For sequencing, libraries were pooled, diluted and sequenced on an Illumina Hi-Seq 2000 using 50bp single-read v3 chemistry. Transcriptome analysis was performed using the Tuxedo suite. TopHat2 was supplied with reads passing vendor quality filtering (PF reads) and the Ensembl transcript set (Homo sapiens, e73, September 2013) as reference. Differential expression was assessed with Cuffdiff v2.1.1. The experiment was performed in triplicate. Fragments per kilo bases of exons for per million mapped reads (FPKM) values and fold changes of significantly up-regulated and down-regulated genes are shown in Supplementary.

### Cell proliferation

LNCaP and PC3 cells were seeded into 6-well plates in triplicate at a density of 10^5^ cells/ml in RPMI 1640 medium (supplemented with 10% FCS). After 24 hours cells were washed with PBS and medium was replaced with Charcoal stripped RPMI 1640 medium (Gibco). 3,3′,5-triiodo-L-thyronine (T3) were purchased from Sigma-Aldrich and dissolved in DMSO at concentrations of 100 mM (T3). Growth medium of LNCaP cells was supplemented with 0.1-100 ng/ml of T3 and the cells were grown for 3 days. PC3 cells were supplemented with 100ng/ml of T3 and were grown for 3 days. After trypsinization and collection by centrifugation, cells were re-suspended and counted using a hemocytometer (C-chip).

### Transfection and transduction of prostate cancer cells

CRYM expressing cells were generated by transfecting PCa cells with pReceiver-M72 Empty Vector and pReceiver-M72 CRYM (for simplicity referred to in the text as CRYM(+) (GeneCopoeia) containing a green fluorescent protein (GFP) reporter gene using Lipofectamine LTX reagent (Invitrogen, Carlsbad, CA, USA) according to the manufacturer’s protocol. GFP expression was evaluated under a fluorescent microscope and cells were selected using 500 μg/mL of G-418 antibiotic for 4 days (Gibco). Cells used for analysis were >70% GFP-positive.

### [^125^I]T3 Uptake experiments

Cell culture medium (RPMI, 10% FCS) used in LNCaP and PC3 cells overexpressing CRYM or empty vector was discarded and replaced with 1 ml of charcoal stripped RPMI per well. Cells were incubated with 45 kBq L-3,5,3’-[^125^I]-triiodothyronine (PerkinElmer, USA) for 12, 24 and 48 hours in an incubator (humidified atmosphere, 37°C, 5% CO_2_). After incubation, the supernatant was taken off the cells. Cells were washed with PBS, trypsinized and centrifuged. Radioactivity was measured in all fractions using a gamma-Counter (2480 WIZARD^2^, PerkinElmer) and uptake values were calculated as percentage of applied dose per 1×10^5^ cells (%AD/ 1×10^5^ cells).

### NMR-Based Metabolomics Analysis of Culture Media and Cell Pellets

The culture media from PC3 were filtrated using 3-kDa cutoff Nanosep centrifugal filters (Pall Life Science) to remove cell debris and macromolecules prior to NMR analysis (Moazzami AA, 2014). In order to remove glycerol from the filter membrane, filters were washed 8 times with 500 ul H_2_O at 36°C shaking at 400xg speed and followed by filtering of 500 μL of culture medium at 4°C at 10000xg. Phosphate buffer (150 μL, 0.4 mol/L, pH 7.0), D_2_O (45 μL) and TSP (30 μL, 5.8 mmol/L) (Cambridge Isotope Laboratories) as internal standard were added to 375 μL of culture media filtrate. The mixture was then used for NMR analysis (5 mm NMR tube). The cell pellet collected from culture media of PC3 was extracted with 1 mL of methanol taken directly from the freezer (−20°C). After centrifugation at 10000xg speed and 4°C, the supernatant was taken and dried using nitrogen. The samples were then dissolved in a mixture containing phosphate buffer (520 μL, 0.135 mol/L), D2O (50 μL) and TSP (30 μL, 5.8 mmol/L) (Cambridge Isotope Laboratories) as internal standard and were analyzed by NMR. The complete sets of NMR quantifications were carried out using a Bruker spectrometer at 600 MHz. ^1^H NMR spectra were obtained using the zgesgp pulse sequence at 25°C, with 200 scans for culture media and 400 scans for cell pellet extract, 65536 data point over a spectral width of 17942.58 Hz. Acquisition duration was selected as 1.82 s, and relaxation delay was chosen as 4.0 s. Bruker Topspin 1.3 software was used to run the NMR spectra and NMR was transformed following multiplication by enlarging of 0.3 Hz. NMR spectra were adjusted to TSP at 0 ppm and baseline together with spectral phase was adjusted. Thirty-nine metabolites on the NMR spectra of the culture media were identified and quantified by manually integrating specific spectral regions after accounting for overlapping signals using NMR Suite 7.1 profiler (ChenomX Inc, Edmonton, Canada). A list of metabolites quantified, their corresponding NMR signals used for integration and quantification, and the order of integration, in order to account for overlapping signals, are presented in Figure S4B. NMR analysis for cell extracts was performed by integrating NMR spectra into 0.01-ppm integral regions (buckets) and residual H_2_O was removed. The identification of NMR signals was carried out using NMR 7.1 library (ChenomX Inc, Edmonton, Canada), Biological Magnetic Resonance Data Bank, and Human Metabolome Data Base.

### Multivariate Data Analysis

The SIMCA-P+ 13.0 software (Umetrics, Umeå, Sweden) was used to perform multivariate data analysis, principal component analysis (PCA) and partial least-squares-discriminant analysis (PLS-DA) as described previously (Moazzami AA, 2011). PCA and PLS-DA models were fitted using metabolomics data from culture media or cell pellet extract of PC3 cells with and without CRYM vector and T3 treatments. In addition, separate PCA models for PC3 cells were fitted using metabolomics data from culture media or cell pellet extract with and without CRYM vector and T3 treatments. For PLS-DA models were fitted using metabolomics data from culture media and cell pellet extract, the variable importance for projection (VIP) for each metabolite or spectral regions were implemented to determine the discriminative metabolites or spectral regions onward the first two components. A metabolites region with VIP ≥ 1 and the corresponding jackknife- based 95% confidence interval (CI) ≤ 0.5 (VIP – CI ≥ 0.5) were considered discriminative. The VIP (CI) of all 39 metabolites (measured in the culture media) along the first and second components of the PLS-DA model are presented in the Figure S4B. For the cell pellet extract, first the spectral regions with VIP – CI ≥ 0.5 were determined and identified (as described above). For those metabolites whose NMR signals appear in a different spectral region, the VIP (CI) of the spectral region with the highest VIP – CI and minimum spectral overlap with other metabolites was chosen for further presentation as relative concentration (Figure S4B).

### FMC-PET/MRI patient study

In the scope of a prospective clinical phase III trial (NCT02659527, EudraCT No.: 2014-004758-33), which was approved by the local Ethics committee (1985/2014) and the national drug authorities, a subgroup of 87 patients out of the preoperative study training cohort was eligible for the presented sub analyses. All patients underwent FMC-PET/MRI for primary staging. Patients gave their written informed consent. Finally, 42 patients underwent radical prostatectomy and post-surgery whole mount sections were available for analyses. The ongoing clinical trial is conducted according to ICH-GCP and the declaration of Helsinki.

### Statistical analyses

After testing for equal distribution of values, a Student’s t-test was used for comparison of groups. If no equal distribution was found we used a Mann Whitney-U test. For calculation of Kaplan-Meier curves we used IBM SPSS Statistics Version 22 with the survival package of R (R Development Core Team, 2009), an open-source language and environment for statistical computing, and *p*-values were calculated using a log-rank test. Statistical methods of survival analyses, as well as follow-up details of our study population have been described in publication (Proestling et al., 2012). We conducted statistical analyses using the R environment v3.5.1 (https://cran.r-project.org).

## Supporting information

Supplemental genes CRYM

## Author Contributions

O.A., O.M., J.P., M.M., G.E., M.H., D.M.H. and L.K. designed the research., O.A., M.S., M.H., J.P., M. Hart., A.A. M., M.Sch., R.Ma., P.A.B., J.B.W., B.M., C.J.R., T.B, J.P., H.A.N.and G.H. performed the experimental work; S.H., S.H. and L.K. performed histological analysis; M.S., G.K. and L.K. provided primary prostate cancer and paraffin tissue specimens; H.A.N. performed Oncomine analyses and prepared figures; O.A., O.M, D.M.H., J.P., S.D.T., M.M., G.H., M.H., Z.C., H.A.N., R.Mo. and L.K. wrote the paper, provided important tools and performed statistical evaluation; respectively.. O.M. M.H. and L.K. contributed to project development wrote manuscript and obtained funding. All authors were involved in critical discussion of the manuscript.

## Acknowledgements

This work was supported by the “Jubiläumsfond der Österreichischen Nationalbank” (grant No. 14856) to O.M.. L.K. was supported by grant FWF, P26011. L.K. and O.M. were supported by the BM Fonds No. 15142 and the Margaretha Hehberger Stiftung No. 15142. J.B.W. was supported by a BBSRC Doctoral Training program (BB/M008770/1). R.M. is supported by the Austrian Science Fund (FWF) (SFB-F04707, SFB-F06105, under the frame of ERA-NET (I 4157-B)). H.A.N. is supported by the FWF, under the frame of ERA PerMed (I 4218-B). R.M. and H.A.N. were also generously supported by a private donation from Liechtenstein. B.M. was supported by a University of Nottingham Vice Chancellors award. D.M.H. was supported by research funds from the University of Nottingham. The CT clinical trial was supported by Siemens and Eckert&Ziegler Radiopharma. We thank Urzula Mc Clurg, Frederic Flamant, Jacques Saramut, Helmut Klocker, Elisabeth Gurnhofer, Georg Hutterer and Sharokh Shariat for important suggestions and support concerning this manuscript.

## Figure legends

**Figure S1.**
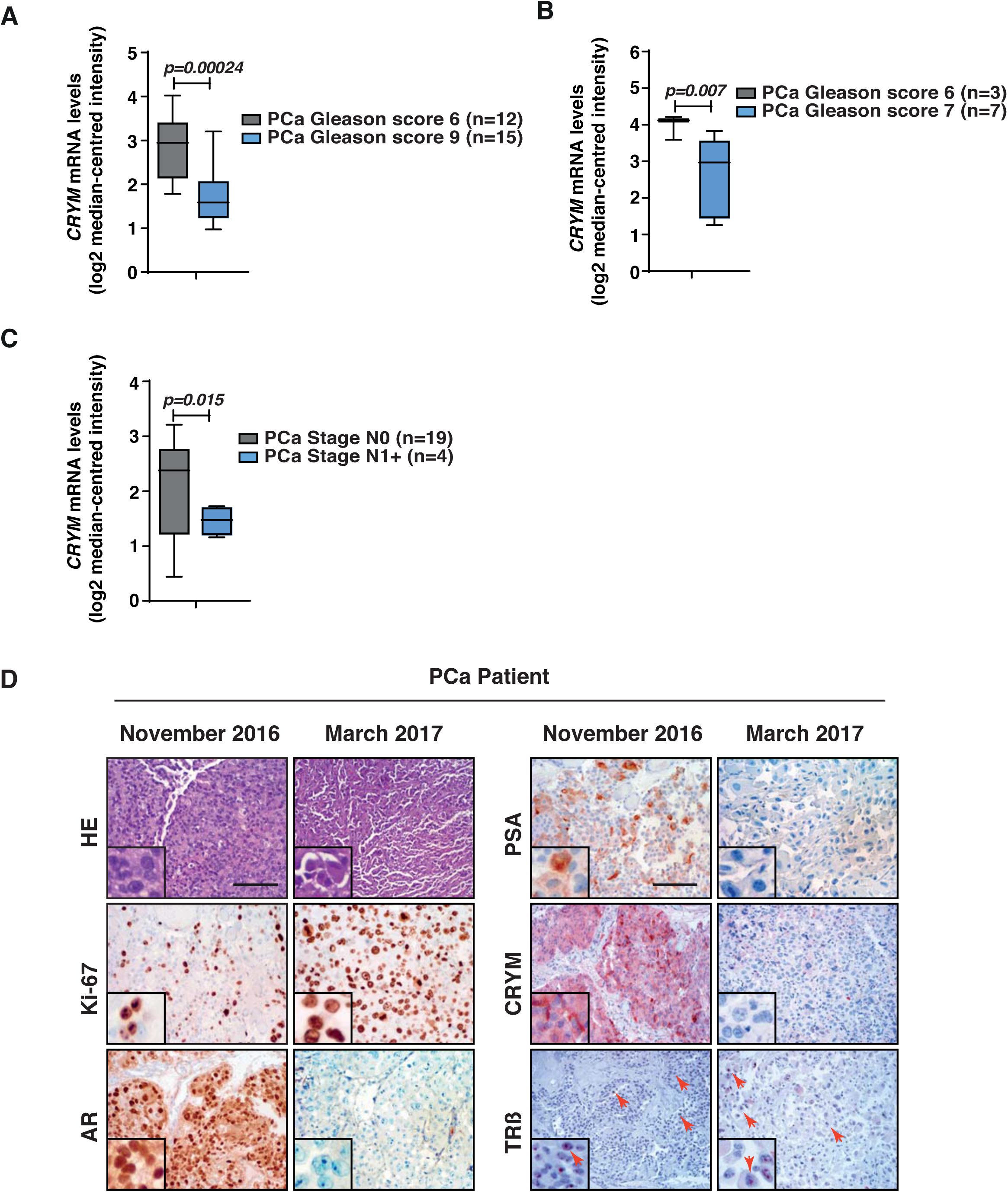
Relative expression of CRYM mRNA. (Log2 median-centered intensity) was compared among PCa subgroups stratified by Gleason score and pathological grade provided from Cancer Microarray Database, Oncomine (Rhodes et al., 2004). **(A)** Relative expression of CRYM mRNA (Log2 median-centered intensity) is compared among subgroups stratified by gleason score. Tumors with Gleason score 9 (n=15) showed low CRYM level in comparison to tumors with Gleason score 6 (n=12). p=0,00024. **(B)** Relative expression of CRYM mRNA (Log2 median-centered intensity) is compared among subgroups stratified by gleason score 6 and 7. Tumors with gleason score 7 (n=7) showed low CRYM level in comparison to tumors with gleason score 6 (n=3). p=0,007. **(C)** Relative expression of CRYM mRNA (Log2 median-centered intensity) is compared among subgroups stratified by pathological grade N0 (n=19) and N1+ (n=4). Tumors with advanced pathological grade N1+ (n=4) showed low CRYM level. **(D)** Longitudinal assessment of tumor progression of a single PCa patient through two consecutive transurethral prostate sections (TURPs) performed on November 2016 and march 2017. Ki67 staining was strongly induced and AR and PSA staining were absent. Remarkably, the patient also lost CRYM expression and TRβ expression was unchanged, suggesting activation of thyroid hormone signaling through loss of thyroid hormone antagonist CRYM.

